# Computational design of two new soluble epoxide hydrolase (sEH) inhibitors

**DOI:** 10.1101/2020.09.17.302026

**Authors:** Jennifer Liem, Sambid Adhikari, Peishan Huang, Justin B. Siegel

## Abstract

Inhibitors of soluble epoxide hydrolase (sEH) enzymes have shown great potential for the treatment of neuropathic pain. However, current sEH inhibitors have poor physicochemical properties and has not been proven to be safe for human treatments yet. New inhibitor designs could have the potential to improve current drugs’ efficacy, and so in this work, chemical intuition and bioisosteric replacement were used to computationally design two novel sEH inhibitors. These new candidates showed good pharmacokinetic properties and presented better docking scores compared to a known sEH inhibitor, t-TUCB, used in the treatment of pain in horses. Homology analysis revealed that *Mus musculus* may not be suitable organism for preclinical trials studies of these novel inhibitors.

## INTRODUCTION

Neuropathic pain is an urgent and unmet clinical issue that can arise from various diseases such as Type I Diabetes^1^ and Alzheimer’s Disease^2^. In the United States alone, the prevalence of neuropathic pain was reported to be around 10% of the population.^3^ Despite causing such high disturbance to patient’s lives, the underlying mechanism of neuropathic pain is poorly understood, and it is hard to treat.

Currently, a major approach to fight neuropathic pain is by inhibiting soluble epoxide hydrolases (sEH), a class of enzymes that convert epoxides into diols. sEH inhibitors (sEHIs) have been shown to exhibit greater potency than non-steroidal anti-inflammatory drugs (NSAIDs).^2^ Most NSAIDs are not effective against neuropathic pain but sEH inhibition was found to have great efficacy against both inflammation and neuropathy.^1,2,4^

There are four known sEHIs that have been studied: Triclocarban (TCC), trans-4-[4-(3-trifluoromethoxyphenyl-l-ureido)-cycloheyloxy]-benzoic acid (t-TUCB), 1-trifluoro-methoxyphenyl-3-(1-propionylpiperidin-4-yl) urea (TPPU) and 1,3-bis(4-methoxybenzyl) urea (MMU). No sEHI has been approved yet for human treatment, but there have been some like t-TUCB tested for treatment in horses.^5,6^ Despite its efficacy, t-TUCB was found to have a low water solubility and high melting point, indicating unfavorable physical properties that leads to poor bioavailability and absorbtion.^7^ Therefore, new drug designs are needed that are more suitable for potential human usage.

The starting point for new designs is the identification of the pharmacophore, a set of functional groups common to all sEH required for inhibition. It has been reported that the general pharmacophore model includes a urea functional group and its activity is enhanced by the presence two large and bulky hydrophobic groups, typically benzene or cyclohexyl rings, on each side of the urea^8^ (Figure 1). The hydrophobic groups are critical as they interact with the hydrophobic pockets of the protein. The inhibition of sEH also depends on hydrogen bonding of the amide group of the inhibitor with the catalytic residue ASP335 of the human sEH. Additionally, the carbonyl of the urea also participates in hydrogen bonding with the residue TYR383.^8,9^

**Figure 1.**
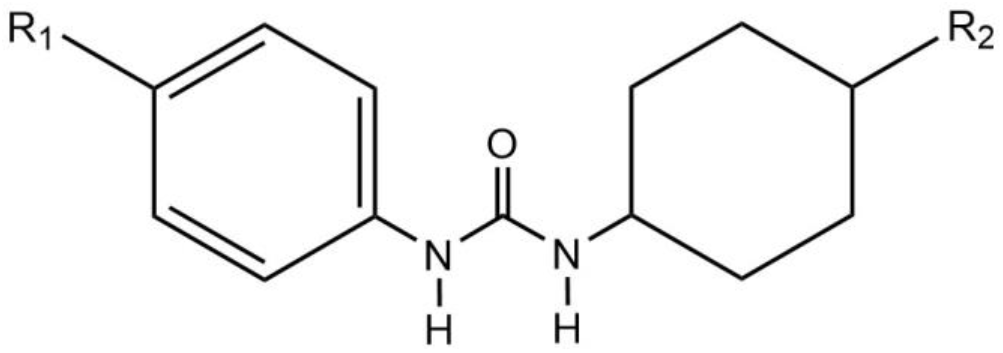
Example of a proposed structure with urea as a central pharmacophore surrounded by two bulky hydrophobic groups attached to an R group.

In this work, keeping the pharmacophore model in mind and using an existing sEHI as a starting point, two novel sEHI drug candidates were designed using computational and chemical intuition-driven methods. Both candidates met the ADMET (Absorption, Distribution, Metabolism, Excretion and Toxicity) criteria and showed improved docking scores compared to current sEHIs. These two new compounds can serve as a possible stepping stone as new treatment for neuropathic pain.

## METHODS

The crystal structure of human soluble epoxide hydrolase complex with t-TUCB (PDB ID: 6AUM) was obtained through the RCSB Protein Data Bank website.^10^

Using the OpenEye software suite, MakeReceptor^11^ was used to define the binding site on the sEH protein. To design a drug candidate based on the pharmacophore using bioisosteric method, vBROOD^12^ was used to generate replacements of functional groups from the model, t-TUCB. SciFinder^13^ was used to confirm that the molecules were not previously studied.

Once the ideal candidates were designed, Gaussview and Gaussian 09^14^ were used to build and optimize drug candidates, respectively. Conformer libraries of known drug molecules and designed candidates were generated using OpenEye OMEGA^15^, after which the conformer libraries were docked into the active site using FRED.^11^

ADMET properties such as hydrogen bond donors and acceptors, molecular weight, number of chiral centers and XLogP bioavailability values were found using OpenEye FILTER.^16^ PyMOL^17^ was used to visualize the protein-ligand inter-actions as well as finding the measurements between the ligand and residues. BLAST^18^ was used to search for human sEH protein homologs. Finally, Jalview^19^ was used to analyze sequences of the sEH protein of various organisms for homology analysis.

## RESULTS AND DISCUSSION

### Evaluation of known Soluble Epoxide Inhibitors

The four known sEHIs, TCC^20^, t-TUCB^5,6^, TPPU^21^ and MMU^22^ were built, optimized and their ADMET properties were analyzed shown in Table 1. None of the inhibitors contained any toxicophore and none were prone to aggregation. Although t-TUCB has a XLogP value higher than five (5.22) which violates Lipinski’s rule, the difference is small and all other properties were found to be favorable, hence, it can still be considered as a starting point. All other molecules passed the Lipinski’s rule of five and were found to be orally bioavailable. The ADMET analysis shows that the known inhibitors have relatively good pharmacokinetic properties.

**Table 1:** ADMET properties and docking results of known inhibitors TCC, t-TUCB, TPPU and MMU

| Name | Molecular Weight (g/mol) | Lipinski H-bond Donor | Lipinski H-bond Acceptor | XLogP | Docking Score |
| --- | --- | --- | --- | --- | --- |
| t-TUCB | 437.39 | 2 | 7 | 5.22 | -18.60 |
| TPPU | 359.34 | 2 | 6 | 2.82 | -16.14 |
| MMU | 300.35 | 2 | 5 | 2.51 | -14.86 |
| TCC | 315.58 | 2 | 3 | 4.53 | -12.40 |

To select one of these sEHI as a starting point for inhibitor design, the four molecules were docked into the human sEH active site and the docking scores were compared (Table 1). TCC was eliminated as a possibility due to its poor score in all four categories. MMU and TPPU’s docking scores were also not as negative as t-TUCB, therefore, with acceptable ADMET properties and possibility of improvement, t-TUCB was selected to be used for drug optimization.

Subsequently, the interactions between t-TUCB and the active site residues of the human sEH was visualized in PyMOL (Figure 2). The main interactions observed are the hydrogen bond between both hydrogens on urea with ASP335 (1.5Å and 1.8Å) and the carbonyl with TYR383 (2.3Å). Additionally, the carbonyl on the carboxylic acid also participates in hydrogen bonding with SER415 (2.5Å). Both phenyl groups on each side of the urea participate in pi-pi T-shaped stacking with TRP525 (3.9Å) and TYR466 (3.9Å) (Figure 2B). By understanding these key interactions, new inhibitors can be designed.

**Figure 2.**
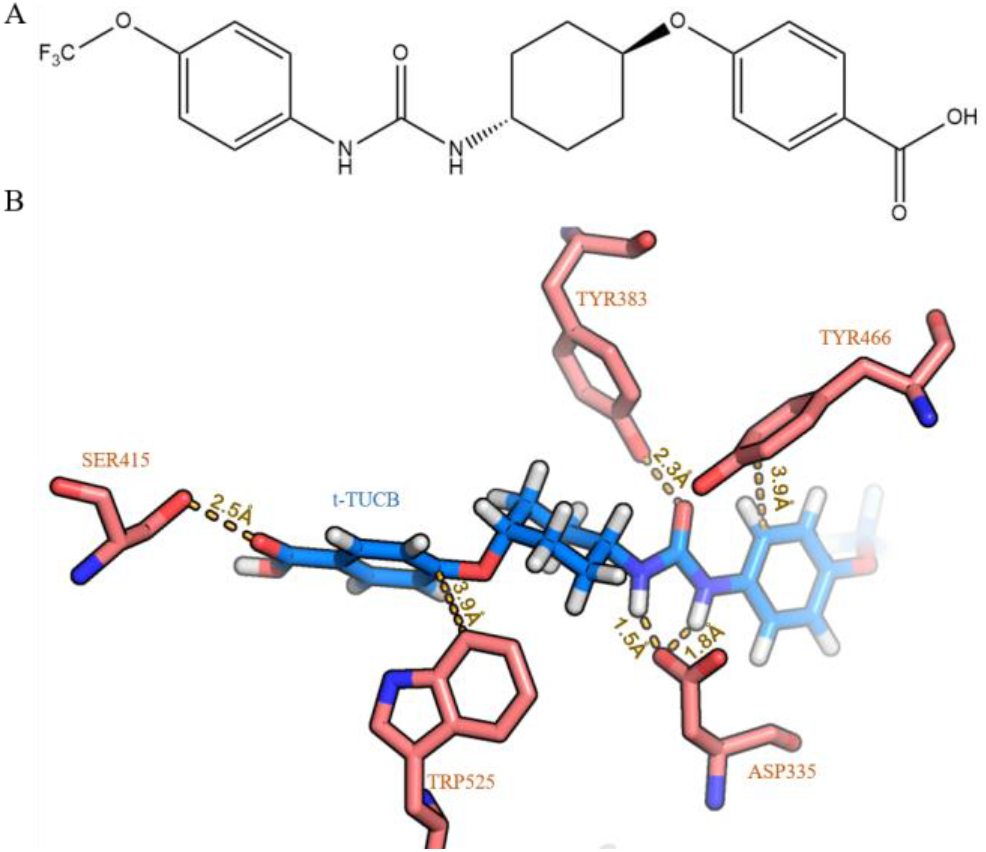
A) 2D Structure of t-TUCB B) t-TUCB (blue) interactions with SER415, TRP525, ASP335, TYR466 and TYR383 residues (pink).

### Computationally Driven Design of Candidate 1

*Candidate 1* was generated by loading t-TUCB onto vBrood 2.0. Based on the docking report, the OCF _3_ group seems to negatively affects the shape score of t-TUCB and does not particularly bind to any residues. Therefore, to improve binding affinity, the OCF_3_ group was chosen for modification. Several compounds that contained the bioisosteric replacement were generated, these were docked into the sEH active site. *Candidate 1* was one of the few bioisosteric replacements that generated a better score. The OCF3 group was replaced with nitrogen and a five membered ring, highlighted in red in figure 3A. As a result, *Candidate 1* has a docking score of −20.80 (Table 2), which is an 11.9% improvement compared to t-TUCB. To rationalize the reason for improvement of the docking score, the interactions between *Candidate 1* and the sEH active site residues in the protein was visualized on PyMol (Figure 3B). The two hydrogen bonds between urea group of candidate 1 with ASP335 were 0.2Å and 0.4Å longer compared to t-TUCB. The hydrogen bond between the carbonyl and TYR383 was shorter by 0.4Å and the hydrogen bond between the carbonyl on the carbolic acid and SER415 is 0.4Å longer.

**Table 2.** ADMET properties and docking results of t-TUCB, *Candidate 1* and *Candidate 2*

| Name | Molecular Weight (g/mol) | Lipinski H-bond Donor | Lipinski H-bond Acceptor | XLogP | Docking Score |
| --- | --- | --- | --- | --- | --- |
| t-TUCB | 437.39 | 2 | 7 | 5.22 | -18.60 |
| *Candidate 1* | 462.56 | 2 | 7 | 5.27 | -20.80 |
| *Candidate 2* | 465.44 | 2 | 7 | 5.88 | -19.57 |

**Figure 3.**
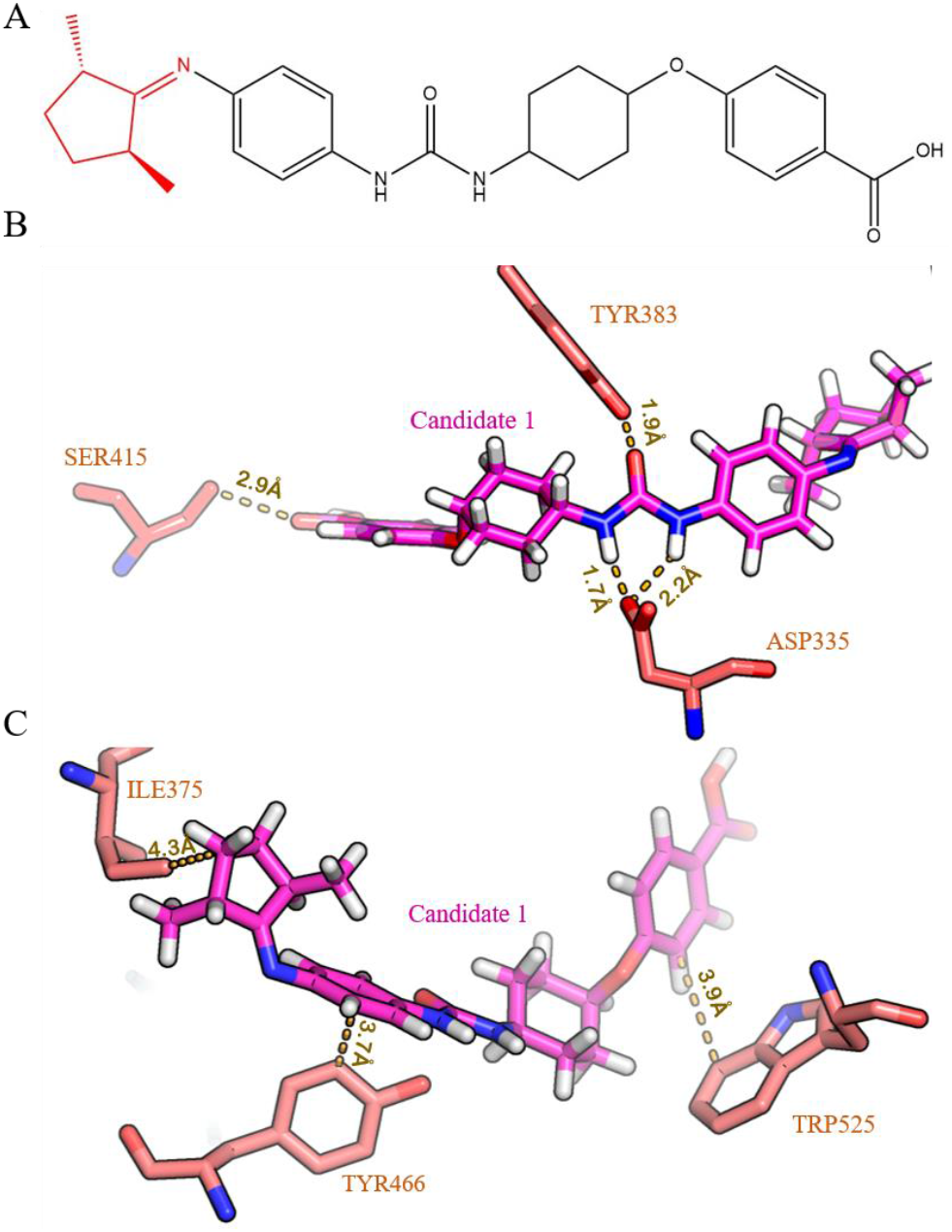
Structure and interactions between Candidate 1 and the human sEH. A) 2D Structure of Candidate 1 with modification highlighted in red. B) Hydrogen bonds between Candidate 1 (fuch-sia) and the active site residues (pink), SER415, TYR383, and ASP335. C) Pi-pi T-shaped stacking between Candidate 1 (fuch-sia) and the active site residues (pink), TRP525 and TYR466 and hydrophobic interaction with ILE375.

*Candidate 1* also retains the pi-pi T-shaped stacking interactions with TRP525 (same bond length of 3.9Å) and with TYR466 (0.2Å reduction in bond length). The main finding in *Candidate 1* is the presence of novel hydrophobic interactions with ILE375 (4.3Å) not present in t-TUCB, due the added ring (Figure 3C). Overall, the new hydrophobic interactions, with the replacement of the new ring structure, can explain the im-provement in docking score of *Candidate 1* compared to t-TUCB.

To further explore whether *Candidate 1* can be administered similarly or better than t-TUCB, its ADMET properties were analyzed using FILTER, and it was found that *Candidate 1* passed the aggregator and filter test (Table 2). Just like with t-TUCB, the XLogP value exceeds 5, which is unfavorable as it affects bioavailability but may be negligible as the increase was not drastic. Based on all the data gathered, *Candidate 1* was found to have suitable pharmacokinetic properties.

### Chemical Intuition Driven Design of Candidate 2

Studies have shown that an important aspect of the pharmacophore are large hydrophobic groups on each side of the urea.^8^ The design of *Candidate 2* was motivated by this fact. In the design of *Candidate 1* above, the addition of the ring enhances binding affinity due to enhanced hydrophobic interactions. Therefore, hypothetically, the addition of an alkyl chain to the benzene ring (Figure 4A) for *Candidate 2* will increase binding affinity similar to the design of *Candiate 1.* Addition-ally, this chain branching will allow the shape of this new candidate to better mimic the natural substrate, epoxy-fatty acids (EpFA).^2^ The docking score (Table 2) of *Candidate 2* is −19.57 which is a 5.2% improvement compared to t-TUCB.

**Figure 4.**
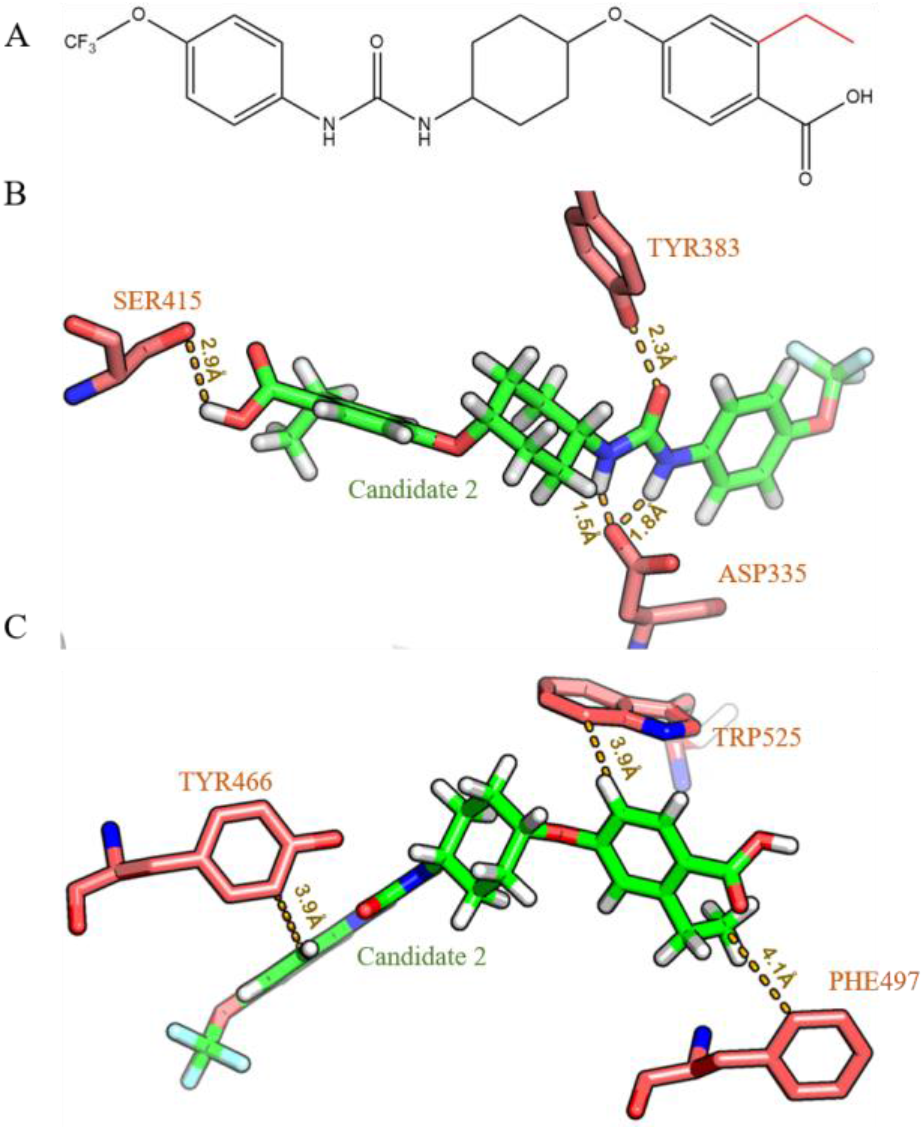
Structures and interactions between *Candidate 2* and the human sEH. A) 2D Structure of *Candidate 2* with modification highlighted in red. B) Hydrogen Bonds between *Candidate 2* (green) and the active site residues (pink), SER415, TYR383, ASP335. C) Pi-pi T-shaped stacking between *Candidate 2* (green) and the active site residues (pink), TRP525 and TYR466 and hydrophobic interaction with PHE497.

To rationalize the improvement in docking score, interactions between *Candidate 2* and sEH was visualized (Figure 4). The hydrogen bond between the carbonyl of the urea and TYR383 and the hydrogens on the urea with ASP335 remained unchanged. The hydrogen bond between the carboxylic acid and SER415 was longer by 0.4Å and the SER415 residue formed a bond with the OH group instead of the carbonyl (Figure 4B). Overall, just like with *Candidate 1*, most hydrogen bonds between *Candidate 2* and active site residues were unchanged or slightly longer for some residues.

Similarly, all existing pi-pi stacking interactions did not change. However, the addition of the alkyl chain leads to new hydrophobic interactions. One such interaction is between PHE497 and the ethyl chain (4.1Å). The new interaction may explain the improvement of the docking score of candidate 2 compared to t-TUCB (Figure 4C).

To further evaluate the potential of *Candidate 2* as a drug, its ADMET properties were generated by FILTER (Table 2). *Candidate 2* also passed both the aggregator and filter test which shows that the compound most likely will not have any potential issues. Unfortunately, XLogP again exceeds 5, which indicates high lipophilicity that may lead to poor absorption but remains roughly similar to t-TUCB and *Candidate 1.* Other than that, *Candidate 2* passed the other criteria for Lipinski’s rule of 5 which suggests oral bioavailability. Although both *Candidate 1* and *Candidate 2* have improved scores, there are potential issues with absorption that might be of concern. However, they are still worth exploring through methods such as animal testing.

### Homolog Analysis

In order to examine the viability of these drug candidates to become potential drugs available in the market for treating neuropathic pain, clinical trials are an important step to test the efficacy. Based on a BLAST search, the results indicated that the soluble epoxide hydrolase of *Mus musculus* and *Oryctolagus cuniculus* are homologs to the human soluble epoxide hydrolase, with percent identities of roughly 73% and 79% respectively. The sequence alignment shows that the human sEH protein conserved the catalytic residue ASP335 (ASP333 in *Mus musculus*) which is critical for inhibition as it forms hydrogen bonds with both hydrogens on the nitrogen of the urea on both candidates.^8,9^ The hydrophobic interacting residue TYR466 (TYR465 in *Mus musculus*) was also conserved in both organisms.

To understand how the candidates interact with the murine sEH receptor compared to humans, the candidates were docked into the murine sEH (PDB ID: 1EK1) ^23^ and their interactions were visualized. The docking score for *Candidate 1* and *2* was −13.83 and −16.48 respectively, both lower than for the human sEH. Interactions between the residues and *Candidate 1* and *Candidate 2* shows that there was no hydrogen bonding with ASP335. Additionally, in both candidates, the pi-pi stacking interaction with TYR465 lost the pi-pi T-shaped stacking with the hydroxyl group of the TYR465 now predicted to point towards the Candidates. These interactions between the residues and *Candidate* 2 is visualized in figure 5. Overall, the modest homology (73%) coupled with the significant energetic and molecular interaction changes observed suggest that *Candidate 1* and *2* will interact differently with *Mus musculus*, limiting its utility as a model organism for preclinical trials for subsequent human clinical trials.

**Figure 5.**
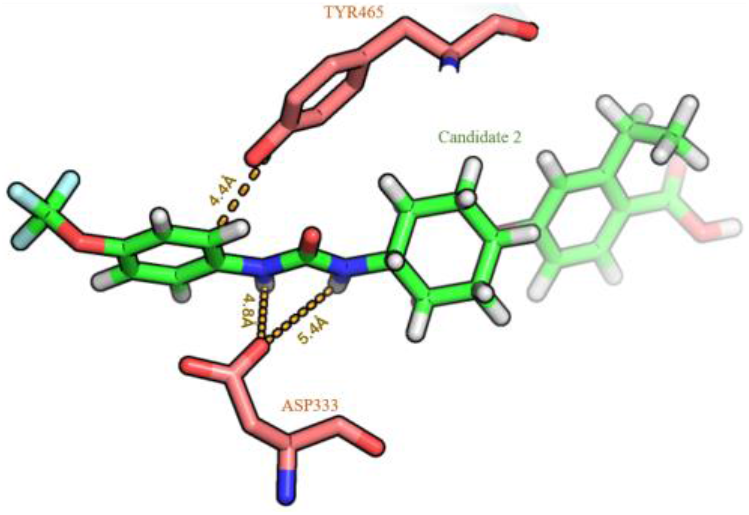
Interactions between *Candidate 2* (green) and the residues (pink) ASP333 and TYR465.

## CONCLUSION

There is a high demand for therapeutics of neuropathic pain as it is almost unbeatable. With the great potential that soluble epoxide hydrolase inhibitors present, it is essential to explore new drug candidates as there are currently none on the market. In this work, two novel drug candidates were designed using computational and chemical intuition-driven methods by modifying t-TUCB.^5,6^ These novel candidates had improved docking scores compared to t-TUCB and showed desirable ADMET properties. Furthermore, homolog analysis was carried out using the *Mus musculus* soluble epoxide hydrolase revealing it may have limited utility for preclinical trials.

Although the data from this computational study shows the candidates exhibited great potential in terms of docking score and similarity in efficacy with t-TUCB, additional analysis and further research is required to properly examine *Candidate 1* and *2*’s ability as a drug and optimize it further, particularly looking to reduce the XLogP scores, improving solubility and lowering the melting point. Furthermore, identification of organisms with higher sEH homology may be critical for both safety and efficacy preclinical evaluation.

## Author Contributions

Research was designed by all authors; all experiments were carried out by J.L. The manuscript was written through contributions of all authors. All authors have given approval to the final version of the manuscript.

## ACKNOWLEDGMENT

Research reported in this publication was supported by UC Davis, the National Science Foundation Award Numbers 1827246, 1805510, 1627539, the National Institute of Environmental Health Sciences of the National Institutes of Health (NIH) under Award Number P42ES004699, UC Davis, NIH Award Number R01 GM 076324-11 and the Rosetta Commons. The content is solely the responsibility of the authors and does not necessarily represent the official views of the National Institutes of Health or National Science Foundation. This study was derived from a course based undergraduate research study conducted in Chemistry 130B at UC Davis.

